# Cellular uptake and viability switch in the properties of lipid-coated carbon quantum dots for potential bioimaging and therapeutics

**DOI:** 10.1101/2024.03.31.587464

**Authors:** Sweny Jain, Nidhi Sahu, Dhiraj Bhatia, Pankaj Yadav

## Abstract

Carbon quantum dots derived from mango leaves exhibited bright red fluorescence. These negatively charged particles underwent coating with the positively charged lipid molecule N-[1-(2,3-dioleyloxy) propyl]-N,N,N-trimethylammonium chloride (DOTMA). However, the bioconjugate displayed reduced uptake compared to the standalone mQDs in cancer cells (SUM 159A), and increased uptake in the case of epithelial (RPE-1) cells. Upon in vitro testing, the bioconjugate demonstrated a mitigating effect on the individual toxicity of both DOTMA and mQDs in SUM-159A (cancerous cells) and of DOTMA in RPE-1 cells. Conversely, it exhibited a proliferative effect on RPE-1 (epithelial cells). Surface modifications of QDs with lipids thus enhances their compatibility with biological systems, reducing systemic toxicity, minimizing off-site effects, sustaining drug release, and modulating cellular viability through various mechanisms (for example, apoptosis), which is, therefore, crucial for multiple applications such as targeted therapeutics.

**TOC:** 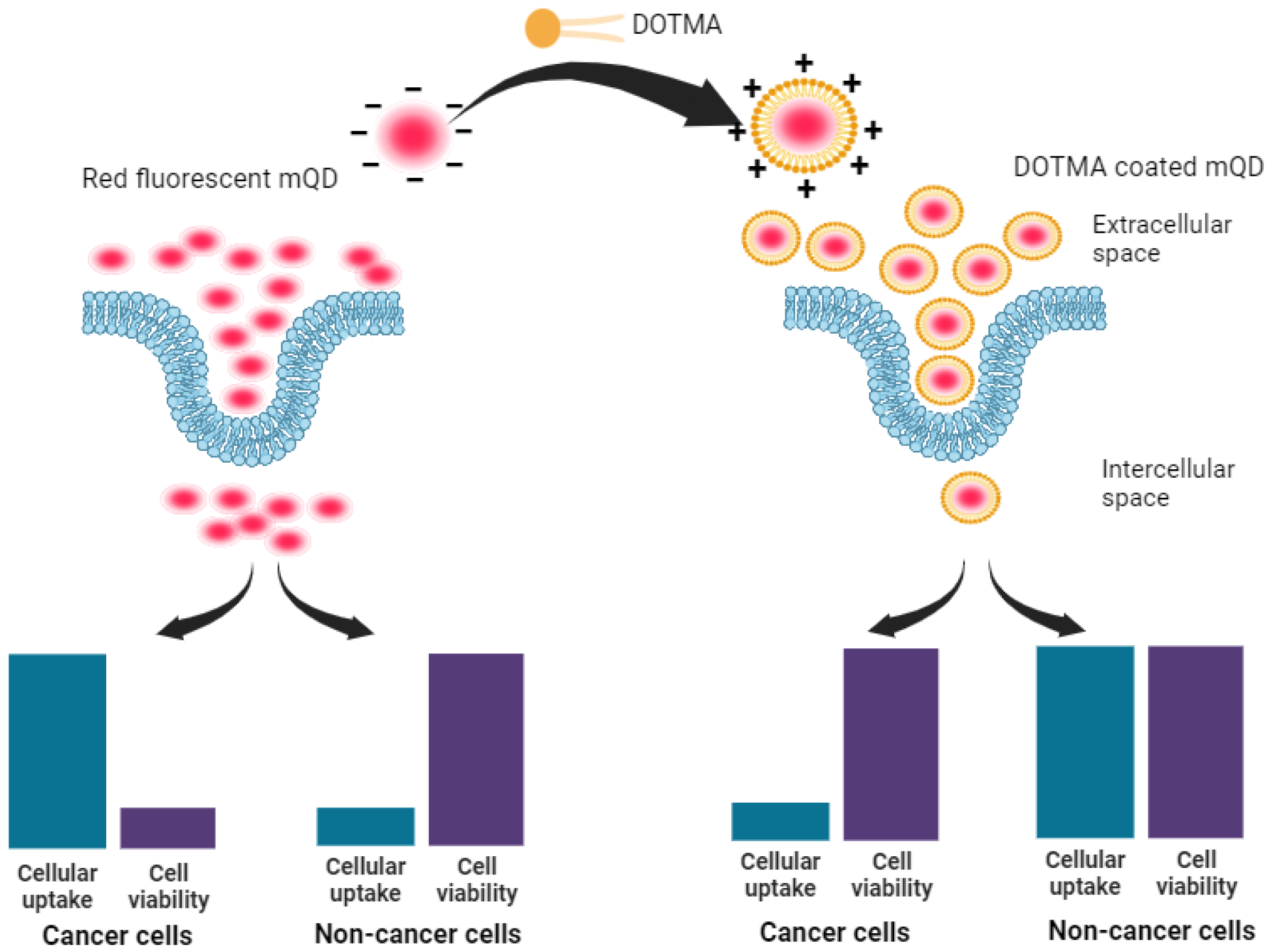

Red emitting, fluorescent carbon quantum dots synthesized using mango leaves(mQDs) showed enhanced cellular uptake and reduced cell viability in the case of cancer cells when compared with lipid-coated mQDs. However, in the case of non-cancerous cells, the lipid-coated mQDs showed enhanced cellular uptake and cell viability when compared with mQDs alone.

## 1. INTRODUCTION

Carbon quantum dots (CQDs) are nanoparticles that exist in zero dimensions, it is a nano-sized semiconductor crystals generally with a size below 10 nanometers^1,2^. Quantum dots (QDs) exhibit distinctive optical characteristics, including a narrow emission peak, a broad excitation spectrum, and robust resistance to photobleaching^3^. These unique optical properties have sparked considerable attention, especially in the realm of biomedical applications^4,5^. It is a type of carbon-based fluorescent nanomaterial. The attributes of CQDs have been harnessed for applications in the field of nanomedicine, specifically in gene therapy, drug delivery, photothermal and radiotherapy, diagnostic techniques, biosensing, the development of fluorescent nanoprobes for bioimaging, and bactericidal activities^6,7^. These materials have attracted growing interest in recent years, primarily because of their exceptional qualities, including environmentally friendly, cost-effectiveness, optical fluorescence with low toxicity, photostability, biocompatibility, extensive surface area, high electrical conductivity, and abundant surface functional groups^8,9^.

Lipids naturally comprise the structural framework of cellular membranes, serving as essential components. Cell membranes are predominantly made up of lipid bilayers, functioning as a protective barrier that shields cellular components from the external environment^10,11^. Lipids have proven to be promising candidates for diverse biomedical applications, Providing a biocompatible protective shield on the surface of nanomaterials^12^. lipids can effectively be used to encapsulate both hydrophilic within the inner aqueous compartment or hydrophobic in the lipid acyl chain region. Indeed, lipids serve as a protective barrier for the enclosed molecular cargo, ensuring its stability^13,14^.

Indeed, lipid molecules can effortlessly diffuse through the membrane, where van der Waals interactions among hydrocarbon chains propel the internalization of the lipid membrane^15^. PEG-lipid and DOTAP-coated SPIOs (Superparamagnetic iron oxide nanoparticles) with an average size of 46nm were investigated for cellular uptake, and biocompatibility and used as a probe for in vivo imaging and the result showed there was an enhancement in the cellular uptake efficiency of cationic-surface L-SPIOs in HeLa, PC-3, and Neuro-2a cells and when Balb/c mice were injected with CT-26 tumor cells loaded with SPIOs, the growth status of these tumors could be tracked using both optical and MR images ^16^.

Poly (lactid-co-glycolid) (PLGA) coated with DOTMA (Lipid-polymer hybrid nanoparticle) with a hydrodynamic size is 250 nm and then mRNA-mCherry was intricately bound with LPNs and then the comparative study was taken for the uptake of fluorescently labeled LPNs loaded with mRNA and An earlier developed polymer-based delivery system (chitosan-PLGA NPs) and it was noticed that LPNs demonstrated a markedly elevated transfection efficiency of approximately 80%, whereas chitosan-PLGA NPs exhibited only around 5%. Translation of the mCherry protein within DCs was also analyzed^17^.

Silica nanoparticles coated with methylene blue (MB) were subsequently double-coated with two lipids—phosphatidylglycerol (PG) and di-oleylphosphatidylcholine (DOPC) with an average size of 117 nm. These coated nanoparticles were employed for comparative investigations of both uncoated and lipid-coated nanoparticles in cellular uptake and localization and analyzing their adsorption on membrane mimics, specifically, Hybrid Bilayer Membranes (HBMs) and it was observed that the adsorption of uncoated nanoparticles to the cellular surface can lead to nanoparticle aggregation, resulting in larger structures compared to silica nanoparticles coated with lipids.^18^

Cationic magnetoliposomes (MLs), characterized by iron oxide cores enveloped in a phospholipid bilayer (dimyristoyl phosphatidylcholine or sphingomyelin) and enriched with cationic lipids (1,2-distearoyl-3-trimethylammonium propane) with a hydrodynamic diameter of 28.8 ± 1.4 nm, are employed as a model to investigate the toxicity associated with cationic lipids and based upon their data they observed that higher doses of cationic lipid may increase electrostatic attraction with the cell membrane, potentially reaching a plateau. This could limit uptake efficiency, regardless of any associated cytotoxic effects^19^.

In recent times, the drug delivery area has shown significant interest in lipid-coated nanoparticles due to their outstanding biocompatibility, effective permeation enhancement, scalability, and broad range of applications. However, the uptake of cationic lipid-coated nanoparticles is influenced by various factors, including the lipid’s properties, surface decoration, size, and physicochemical attributes of liposome formulations. These elements collectively impact the potential release of cationic lipids from the nanoparticle and its subsequent movement toward the cell plasma membrane^19,20^.

We introduce novel red-emitting carbon dots, mQDs, derived from mango leaves, with an emission wavelength of 670 nm and a size of 10.1 nm. These mQDs were decorated with a small cationic lipid, N-[1-(2,3-dioleyloxy)propyl]-N,N,N-trimethylammonium chloride (DOTMA), forming a bioconjugate mQDs: DOTMA, which exhibited enhanced fluorescence intensity, photostability, and reduced cytotoxicity compared to standalone mQDs and DOTMA. Our findings highlight the bioconjugate’s potential for bioimaging, offering brighter signals and improved biocompatibility for precise cellular imaging in *invitro* studies. Cellular internalization assays and viability assessments revealed decreased uptake in cancer (SUM159A) and increased uptake in retinal pigment epithelial (RPE-1) cells, demonstrating the bioconjugate’s promising utility in cellular imaging and potential therapeutic applications.

## 2. Materials and Methods

### 2.1. Materials

Mango leaves were obtained from mango trees at the IIT Gandhinagar campus. Silicon oil was purchased from X-chemicals, ethanol (>99.9%) from Changshu Hongsheng Fine Chemicals Co. Ltd. The filter of 0.22 micrometer was purchased from Merck, as well as the deionized water was taken from Merck Millipore. Cationic lipid, *N*-[1-(2,3-dioleyloxy)propyl]-*N,N,N*-trimethylammonium chloride (DOTMA), was purchased from Avanti Polar Lipids. Phalloidin was purchased from Sigma Aldrich. The cell culture dishes, dimethyl sulfoxide (DMSO), and rhodamine B were purchased from Himedia. DMEM (Dulbecco’s modified Eagle’s medium), Ham’s F12 media, FBS (Fetal bovine serum), and trypsin-EDTA (0.25%) were obtained from Gibco. All the purchased chemicals were of analytical grade without the need for further purification.

### 2.2. Methods

#### 2.2.1 Synthesis of mQDs

The mango leaves were plucked from a mango tree on the IIT Gandhinagar campus. Later, these leaves were washed and dried at ambient temperature for ten days. These dried leaves were then ground to form a fine powder. Next, mango leaves and ethanol are taken in a ratio of 1:10 (w/v) and kept for stirring for 4 hours. This mixture is then centrifuged at 10,000 rpm for 10 minutes at 25°C. The supernatant is then refluxed for 2 hours at 160°C. Once the solution is cooled, mQDs are thus obtained. Next, the solvent is removed using a rotary evaporator. This is further characterized and functionalized with DOTMA to study its effects *in vitro*.

#### 2.2.2. Characterization of mQDs

mQDs were analyzed for optical properties using both UV-Vis absorbance and fluorescence emission using Spectrocord-210 Plus Analytokjena (Germany) and FP-8300 Jasco spectrophotometer (Japan) respectively. For AFM sample preparation mica sheets were peeled freshly and 5μL of samples (1:0.25-1:10 were dropped having concentration (100ug/mL). The sheet containing samples was then placed in a desiccator for drying overnight. Finally, the AFM imaging was done in tapping mode using the Bruker AFM instrument. FTIR spectra of all the concentrations of mQD:DOTMA were recorded from 450 cm^−1^ to 4000 cm^−1^ using spectrum 2 PerkinElmer in ATR mode. Further, analysis of the powder was done for XRD diffraction pattern using Bruker-D8 Discover with a speed of 0.2/min from 5°to 90°. The Malvern analytical Zetasizer Nano ZS was utilized to conduct Dynamic Light Scattering (DLS) for the solution-based size characterization of mQDs and mQDs: DOTMA. Subsequently, the obtained data was plotted using Gaussian fit in the OriginPro software. The photostability of mQDs was analyzed by incubating the samples for 10 days. Fluorescence intensity readings at 670 nm were taken every day following excitation with 400 nm light. The relative fluorescence intensity was plotted by normalizing each fluorescence intensity measurement against the maximum fluorescence intensity recorded. These mQDs and mQD:DOTMA conjugates were then dissolved in the serum-free media to assess the cellular uptake and cellular uptake of these particles in the breast cancer cell line (SUM-159A), and healthy cells (RPE-1).

#### 2.2.3. Cell culture and cellular uptake assay

For the cellular uptake experiment, the RPE1 cells were cultured in DMEM, and SUM-159A (breast cancer) cells were maintained in HAMS-F12 media containing 10% fetal bovine serum and antibiotic at 37 °C with 5% CO2 in a humidified incubator. Approximately 10^5^ per well cell counts were seeded on a glass coverslip in a 24-well plate overnight. Before treatment, the seeded cells were washed with 1× PBS buffer three times and then incubated in serum-free media for 15 min at 37 °C with 5% CO2 in a humidified incubator. After washing, the cells were treated with mQD and mQD:DOTMA to assess their cellular internalization. Different combinations of mQD–DOTMA (mQD:DOTMA; 1 : 0.25, 1 : 0.5, 1 : 1, and 1 : 2, 1:5. 1:10) were used for cell treatment. The treated cells were fixed for 15 min at 37 °C with 4% paraformaldehyde and rinsed three times with 1×PBS. The cells were then permeabilized with 0.1% Triton-X100 and stained with 0.1% phalloidin to visualize the actin filaments. Then the cells were washed three times with 1× PBS and mounted onto the slides with Mowiol and DAPI to stain the nucleus.

#### 2.2.4. MTT Assay

An MTT assay was performed to assess the effect of cytotoxicity of the synthesized mQD-DOTMA. Cells were seeded in 96-well plates at a seeding density of 5000 cells per well. The culture plates were incubated at 37 °C for 24 h. The cells were treated with different ratios of mQD–DOTMA (1:0.5, 1 : 1, 1:2, 1 : 5, 1 : 10) at varied concentrations (100ug/mL, 200ug/mL, 300ug/mL). Then they were incubated at 37 °C for 24 h. Untreated cells served as a control. After incubation, 0.5 mg ml^−1^ of a 3-(4,5-dimethylthiazol-2-yl)-2,5-diphenyltetrazolium bromide (MTT) solution was added to each well and incubated at 37 °C for 4 h. The solution was removed and replaced with dimethyl sulfoxide (DMSO) in each well and incubated in the dark for 15 min to dissolve the formazan crystal. A multiwell microplate reader was used to measure absorbance at 570 nm. The experiment was conducted in triplicate, with normalization to the corresponding well containing DMSO. The non-treated mQDs well served as the control to determine the % cell viability of each well. The cell viability percentage was calculated using the following formula:

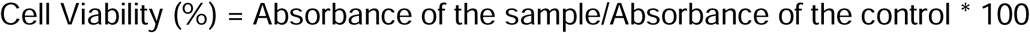

#### 2.2.5. Confocal Imaging

Confocal imaging was conducted on fixed cells (using a 63x oil immersion objective) and fixed tissues/embryos (using a 10x objective) employing the Leica TCS SP8 confocal laser scanning microscope (CLSM) from Leica Microsystems, Germany. Various fluorophores were excited using different lasers: DAPI (405 nm), and mQDs (633 nm). The pinhole aperture was maintained at 1 airy unit throughout the imaging process. Quantitative analysis of the images was carried out using Fiji ImageJ software. The analysis involved measuring the whole cell intensity from maximum intensity projections, subtracting background, and normalizing the measured fluorescence intensity against unlabeled cells. Approximately 40-50 cells were quantified from the collected z-stacks for each experimental condition.

#### 2.2.6. Statistical Analysis

GraphPad Prism software (version 8.0.2) was employed for statistical analysis. Data were presented as means ± standard deviation (SD) or means ± standard error from two independent experiments. p values were computed using one-way ANOVA and two-tailed unpaired Student t-tests, with a 95% confidence interval.

## 3. Results

### 3.1. Characterization of mQDs and mQDs:DOTMA

Red emitting, fluorescent carbon quantum dots were obtained using mango leaf powder. Freshly plucked mango leaves were dried in the room light. The dried mango leaves were stirred with ethanol in 1:10 (weight/volume) ratio. The supernatant is then refluxed at 160°C for 2 hours and then filtered using 0.22μm filter. The quantum dots are formed. To obtain a powder of it rotavapor was done as illustrated in **Figure 2**.

**Figure 1.**
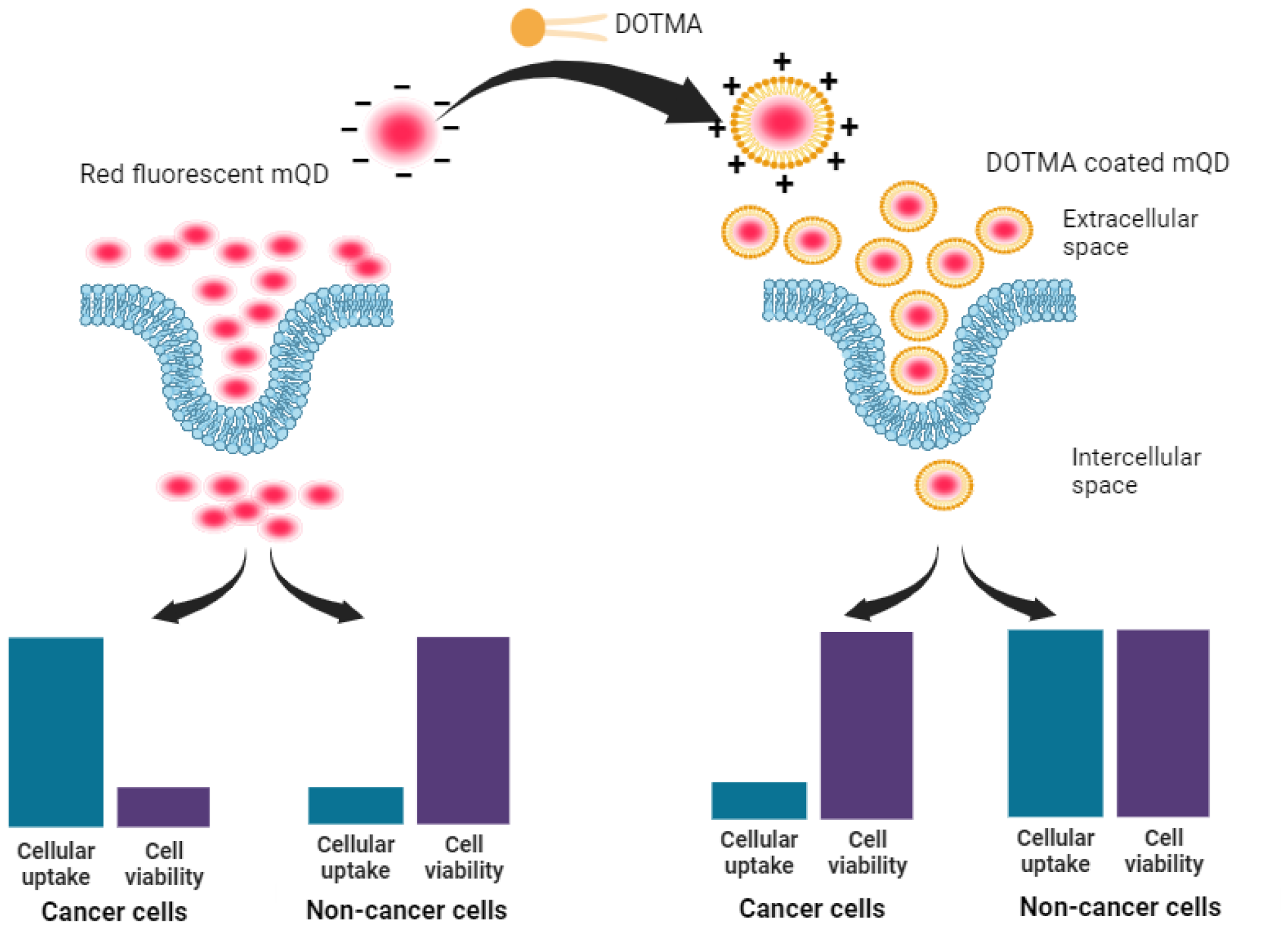
mQDs coated with DOTMA increase the cell viability in both cancerous (SUM-159A) and non–cancerous (RPE-1) cells, However, the uptake is switched from higher to lower in cancer cells and from lower to higher in non-cancerous cells, which can be used for biomedical applications.

**Figure 2:**
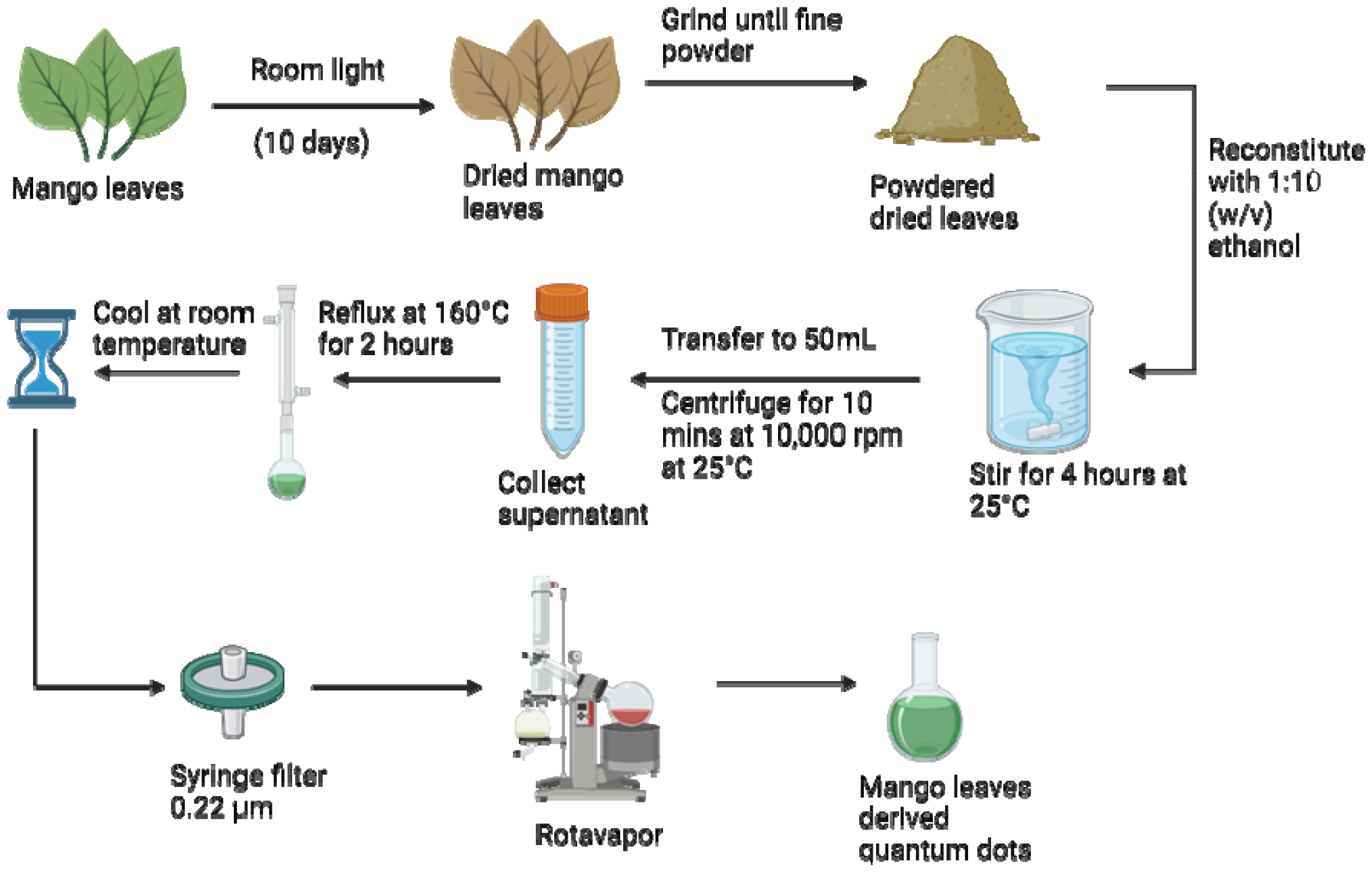
Synthesis of carbon quantum dots (mQDs) using mango leaves as the precursor material. The fresh mango leaves were dried at room temperature and then powdered. The powdered mango leaves were dissolved in ethanol and centrifuged after 4 hours to collect mango leaf extract. The extract was then subjected to reflux at 160 degrees Celsius for 2 hours and then syringe filtered after cooling down to room temperature. The solvent is then evaporated using rotavapor to get the powdered mQDs.

The mQDs thus formed has an inherent negative charge, this prompted us to conjugate it electrostatically with the positively charged small lipid molecule, N-[1-(2,3-dioleyloxy)propyl]-N,N,N-trimethylammonium chloride (DOTMA). The mQDs and DOTMA were mixed in the following ratios: 1:0.25, 1:0.5, 1:1, 1:2, 1:5 and 1:10 with water as the solvent. Next characterization studies were carried out for mQD and the bioconjugates. To characterize the quantum dots thus formed the optical properties were analyzed using UV Spectra and Fluorescence Spectra. In UV spectra, for DOTMA there were no peaks observed, whereas peaks were observed in case of mQDs at 206nm (-OH group), 260nm (flavonoids) and the mQD: DOTMA conjugate displayed both the peaks of mQDs and another peak at 327nm as shown in **Figure 3(A)**. Further, for fluorescence spectra 400nm wavelength was chosen as it gave the maximum intensity. Fluorescence spectra of DOTMA showed a peak at 460nm, and both mQD and mQD:DOTMA conjugate displayed peaks at 460nm, 495nm, 549nm and 679nm upon excitation at 400nm as shown in **Figure 3(B)**. As the DOTMA concentration increased the fluorescence intensity of the sample increased **(Figure 3(B))**. Hence, we observed presence of DOTMA not only enhanced the fluorescence of mQD but also increased the photostability of mQDs **(Supplementary Figure 1)**. Next to understand the functional groups present on our quantum dots we performed FTIR, and the spectra show the presence of different functional groups present on the surface of mQD: 2923 cm-1 for alkanes and acidic groups; 2850 cm-1 for aldehyde, alkanes, acidic groups; 1560 cm-1 for nitro compounds; 1188 cm-1 for anhydride (acyl), amines, alcohol (alkoxy), esters; 938 cm-1 for alkenes (sp2-C-H)as shown in **Figure 3(C)**.

**Figure 3:**
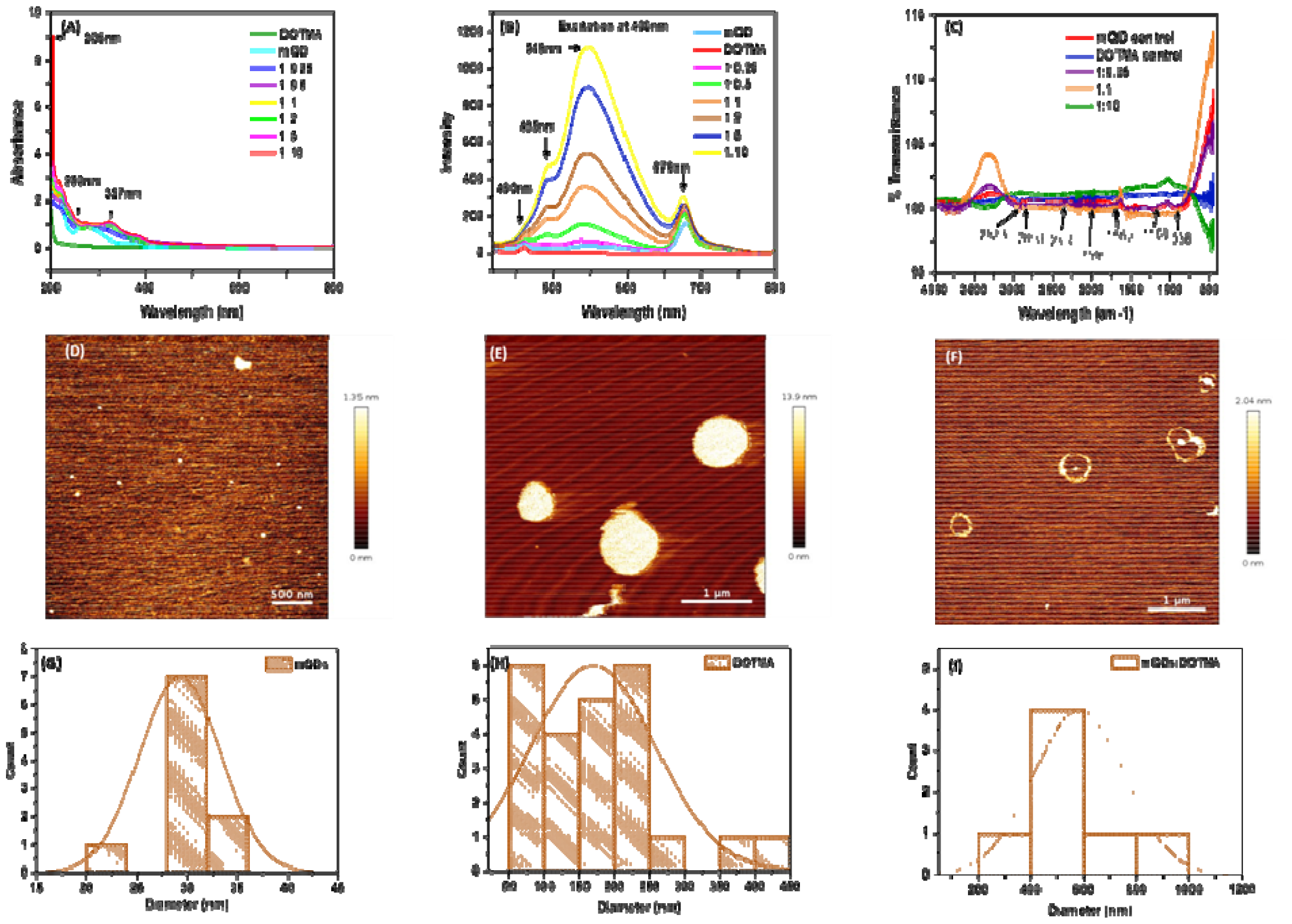
Characterization of the conjugate formed of mQD and DOTMA. (A) The UV spectra of the DOTMA showed no peaks, whereas peaks were observed in case of mQDS at 206nm (-OH group), 260nm (flavonoids) and the mQD: DOTMA conjugate displayed both the peaks of mQDs and another peak at 327nm. (B) Fluorescence Spectra of DOTMA showed a peak at 460nm, and mQD and mQD:DOTMA conjugate displayed peaks at 460nm, 495nm, 549nm and 679nm upon excitation at 400nm. (C) The FTIR spectra show the presence of different functional groups present on the surface of mQD: 2923 cm-1 for alkanes and acidic groups; 2850 cm-1 for aldehyde, alkanes, acidic groups; 1560 cm-1 for nitro compounds; 1188 cm-1 for anhydride (acyl), amines, alcohol (alkoxy), esters; 938 cm-1 for alkenes (sp2-C-H). (D)The quasi-spherical morphology and topography of mQDs were studied using atomic force microscopy (AFM). (E) The quasi-spherical morphology and topography of DOTMA were studied using atomic force microscopy (AFM). (F) The ring-like structure of DOTMA coating around the mQD can be observed using atomic force microscopy (AFM). (G) Histogram plot of AFM analysis of mQD showing their size at around 29nm. (H) Histogram plot of AFM analysis of DOTMA showing their size at around 173nm. (I) Histogram plot of AFM analysis of mQD:DOTMA conjugate showing their size around 580nm.

To analyze the size and morphology of the quantum dots Atomic Force Microscopy (AFM) and Dynamic Light Scattering (DLS) was performed for both mQD and mQD:DOTMA conjugates. The quasi-spherical morphology and topography of mQDs, DOTMA were observed with a size of about 30nm and 179nm, respectively (**Figure 3(D,E,G,H)**). The ring-like structure of DOTMA coating around the mQD can be observed in case of conjugate which was about 580nm (**Figure 3(F) and (I)**).

mQDs and DOTMA were mixed in water as a solvent and the following ratios were made: 1:0.25, 1:0.5, 1:0.75, 1:1, 1:2, 1:5, 1:10. The hydrodynamic size and zeta potential were assessed following the conjugation of mQDs with DOTMA as shown in **Figure 4 (A-I)**. The hydrodynamic size of mQDs and DOTMA alone in water as solvent was measured as 10.1 nm and 142 nm, respectively. The hydrodynamic radius of mQDs: DOTMA (1:0.25, 1:0.5, 1:0.75, 1:1, 1:2, 1:5, 1:10) in water was measured as 122nm, 122nm, 122nm, 122nm, 164nm, 190nm, 190nm respectively. The size of the conjugates 1:0.25, 1:0.5, 1:0.75 and 1:1 showed a similar size as the mQDs surface is not yet saturated with the lipid layer of DOTMA and are still forming the lipid layer/coating. As the concentration increases further the size of the bioconjugate increases with increase in concentration of DOTMA as new lipid layer is formed. A similar trend is observed in 1:2, 1:5 and 1:10 ratios where the size of the bioconjugate is approximately 164nm, 190nm and 190nm as the surface gets saturated with DOTMA. Another saturation is observed in ratios, 1:5 and 1:10, as even on increasing the concentration double the amount the hydrodynamic radius remained same. The Zeta Potential of our negatively charged molecule, mQDs is –0.99mV and that of positively charged DOTMA is 21.1mV. We can observe an increase in the positive charge in the conjugates as the concentration of DOTMA increases as shown in **Figure 4 (J)**.

**Figure 4:**
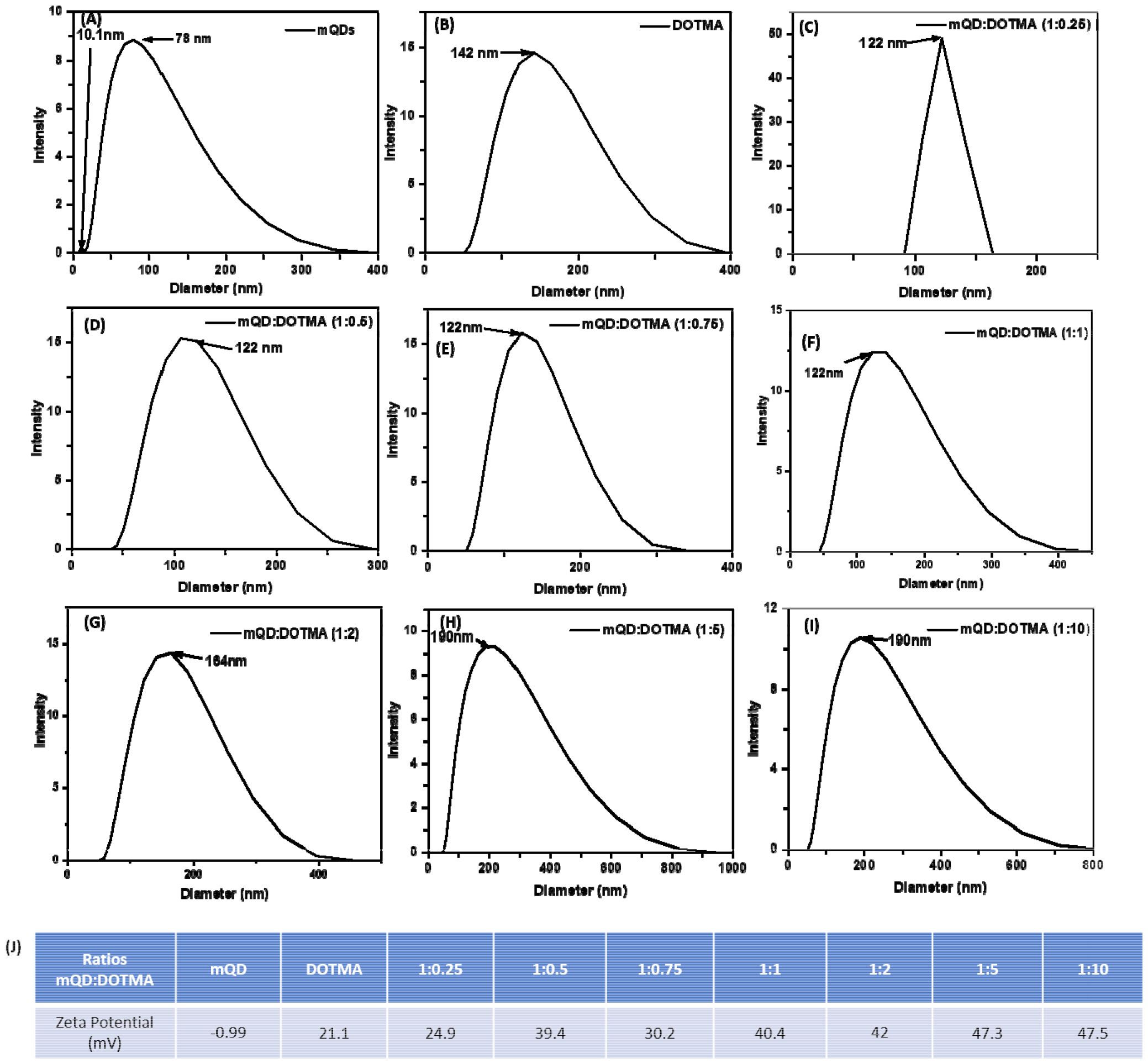
Studying hydrodynamic radius and charge using Dynamic Light Scattering. (A) the hydrodynamic radius of mQD in water as solvent has two sizes 10.1nm and 78nm. (B) the hydrodynamic radius of DOTMA in water is 142nm. (C) The hydrodynamic radius of 1:0.25 ratio of mQD:DOTMA conjugate is 122nm. (D) The hydrodynamic radius of 1:0.5 of mQD:DOTMA conjugate is 122nm. (E) The hydrodynamic radius of 1:0.75 of mQD:DOTMA conjugate is 122nm. (F) The hydrodynamic radius of 1:1 of mQD:DOTMA conjugate is 122nm. (G) The hydrodynamic radius of 1:2 of mQD:DOTMA conjugate is 164nm. (H) The hydrodynamic radius of 1:5 of mQD:DOTMA conjugate is 190nm. (I) The hydrodynamic radius of 1:10 of mQD:DOTMA conjugate is 190nm. (J) The Zeta Potential of mQDs, DOTMA and their ratios (in mV).

### 3.2. Cytotoxic study of mQDs and mQDs:DOTMA

To investigate the cell-specific responses to mQDs and mQD:DOTMA, we conducted 3-[4,5-dimethylthiazol-2-yl]-2,5-diphenyltetrazolium bromide (MTT) assays on RPE1 epithelial cells and SUM159A breast cancer cells. The assay was done with increasing concentrations of mQDs (100, 200, 300μg/mL) and ratios (1:0.5, 1:1, 1:2, 1:5, 1:10) of mQD:DOTMA (100, 200, 300μg/mL) were administered to the cells. After a 24-hour incubation period, we calculated the percentage viability for both the cell types.

In SUM-159A cells we observed with mQDs alone at concentrations 100, 200, 300μg/mL showed cell viability of 12%, 16%, 33% respectively. For DOTMA alone at concentration 100, 200, and 300μg/mL is 18%, 8%, 8% respectively. This established that in SUM-159 cells alone mQD and alone DOTMA are toxic to the cells hence the cell viability has decreased significantly. For ratios 1:0.5, 1:1, 1:2, 1:5 and 1:10 concentrations 100, 200 and 300μg/mL were taken. The cell viability for ratio 1:0.5 is 38%, 104%, 104% respectively. The cell viability for ratio 1:1 is 64%, 53%, 68% respectively. The cell viability for 1:2 is 23%, 47%, 68% respectively. The cell viability for 1:5 is 26%, 30%, 27% respectively. And for ratio 1:10 cell viability is 38%, 22%, 21% respectively. Hence, we can see that 1:0.5 ratio of mQD:DOTMA at 200 and 300μg/mL concentration is not toxic to the cell and can be further explored as a potential molecule for bioimaging purposes.

Further with respect to RPE cells, mQDs alone at 100, 200, 300μg/mL showed cell viability of 71%, 80%, 60%. For DOTMA alone the cell viability is 33%, 37%, 42%. These findings suggest that DOTMA when given alone to the cells is toxic. For ratios 1:0.5, 1:1, 1:2, 1:5 and 1:10 concentrations 100, 200 and 300μg/mL were taken. The cell viability for ratio 1:0.5 is 150%, 252%, 292% respectively. The cell viability for ratio 1:1 is 226%, 284%, 370% respectively. The cell viability for 1:2 is 227%, 315%, 353% respectively. The cell viability for 1:5 is 83%, 111%, 164% respectively. And for ratio 1:10 cell viability is 123%, 139%, 92% respectively. We can see that when mQDs are conjugated with DOTMA it quenches the cytotoxic effect and further leads to cell proliferation as seen in **Figure 5**.

**Figure 5:**
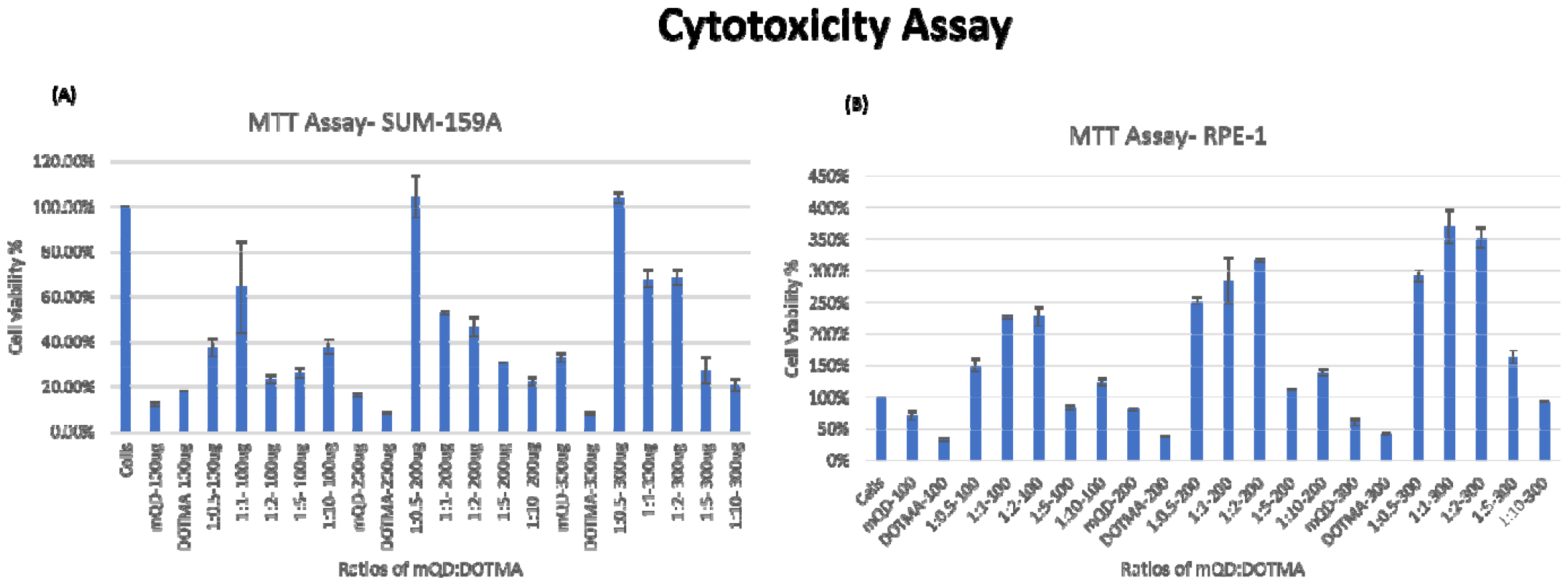
Cytotoxicity Assay **(A)** Treatment of different concentrations (100, 200, and 300μg/mL) of mQD, DOTMA, and mQD: DOTMA biconjugate in SUM-159A. The mQDs and DOTMA alone at all the concentrations (100, 200, and 300μg/mL) showed cell viability of less than 20% except in the case of mQDs 300μg/mL of around 30%. However, the lipid-conjugated mQDs showed enhanced cellular viability as compared to mQDs and DOTMA alone. The 1:0.5 bioconjugate (mQDs: DOTMA) showed enhanced cellular viability of 60% in case of 100μg/mL, and more than 100% in case of 200μg/mL and 300μg/mL, but the rest of the ratios in all concentrations showed less than 60% cell viability. **(B)** Treatment of different concentrations (100, 200, and 300μg/mL) of mQD, DOTMA, and mQD: DOTMA biconjugate in RPE-1 cells. The mQDs at all the concentrations (100, 200, and 300μg/mL) showed cell viability of around 60-70%, and DOTMA alone at all the concentrations (100, 200, and 300μg/mL) showed cell viability of less than 30%. However, the lipid-conjugated mQDs showed enhanced cellular viability as compared to mQDs and DOTMA alone. The 1:0.5, 1:1, 1:2 bioconjugate (mQDs: DOTMA) showed enhanced cellular viability of more than 150% in the case of all the concentrations (100, 200, and 300μg/mL) having 350% in case of 1:1 of 300μg/mL. However, the cell viability reduced back to 100% in 1:5 and 1:10 bioconjugate (mQDs: DOTMA) in all the concentrations (100, 200, and 300μg/mL) except 1:10 bioconjugate (mQDs: DOTMA) in 200μg/mL and 1:5 bioconjugate (mQDs: DOTMA) in 300μg/mL having 150% cell viability.

### 3.3. Cellular Uptake of mQDs and mQDs: DOTMA

After characterization, these mQDs and mQD: DOTMA conjugates were employed to examine their impact on cell lines. Their effects were assessed on SUM-159A cancerous cells, considering previous reports indicating the anti-cancer properties of mQDs. Fluorescence signals emitted by mQDs, alongside standard markers like DAPI (for the nucleus), aided in visualizing cellular morphology and physiology. Based on the toxicity studies, two experiments were designed to validate our results. Two types of cells were taken for the study, namely cancer cells (SUM159A) and non-cancerous epithelial cells (RPE-1). The cells were seeded and later treated with mQDs and mQDs: DOTMA to observe the effect.

In the case of both cancer cells and epithelial cells, there was an increase in uptake at concentration 100 and 200 μg/mL of mQDs compared to 50 μg/mL and control. Thus, the cellular uptake studies were performed with 100μg/mL and 200μg/mL concentrations as shown in **Figure 6**.

**Figure 6:**
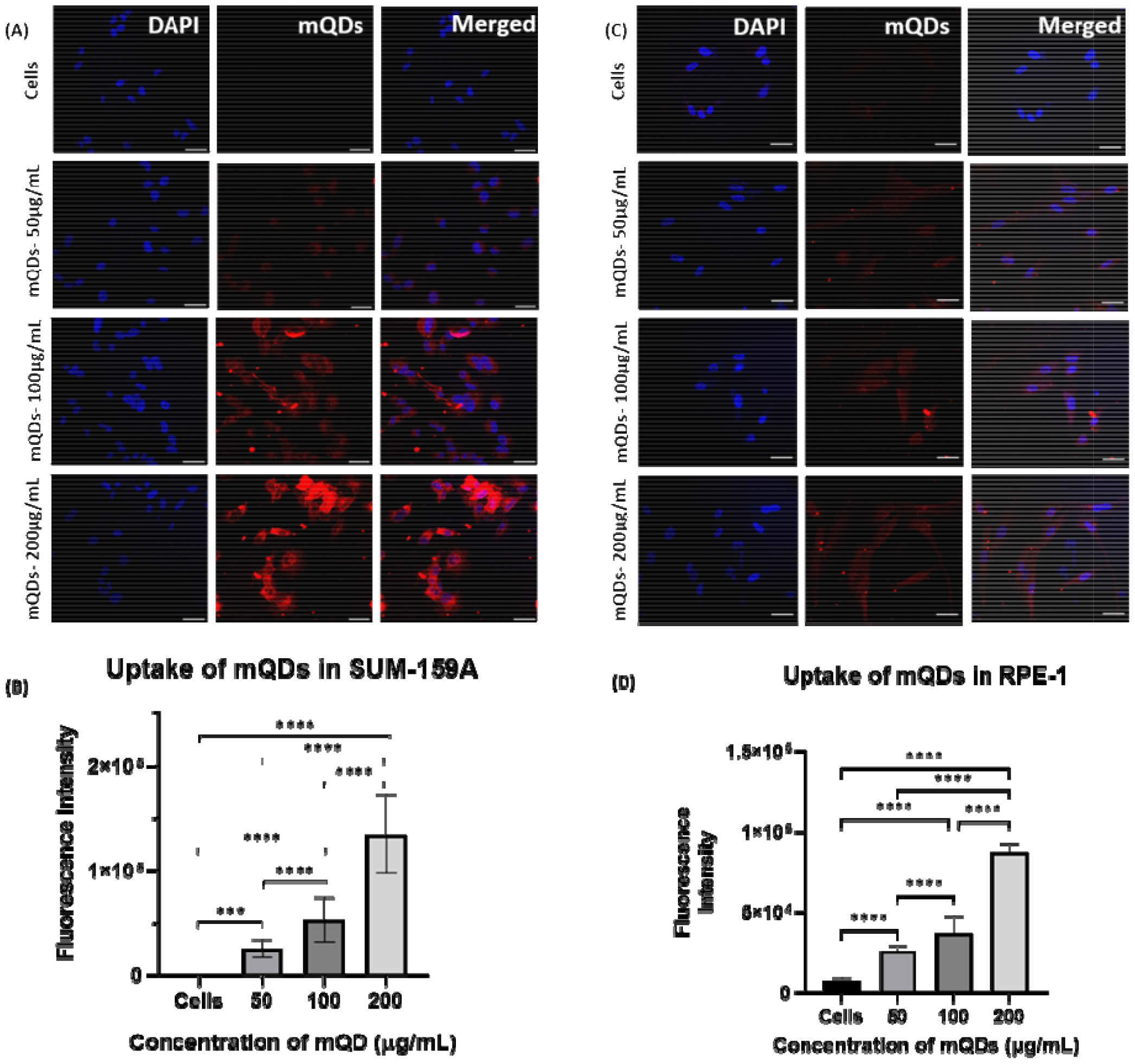
Cellular uptake of mQDs in SUM-159A and RPE-1. Scale bar-5μm (A) Uptake of mQDs at concentrations 50μg/mL, 100μg/mL, 200μg/mL in SUM-159A. (B) Quantified fluorescence intensity of mQDs in SUM-159A. (C) Uptake of mQDs at concentrations 50μg/mL, 100μg/mL, 200μg/mL in RPE-1. (D) Quantified fluorescence intensity of mQDs in RPE-1. The statistical significance was tested by one-way ANOVA in the Prism Software and is represented as **** when p < 0.0001 and ns when there is no significant difference. (n = 30 cells per condition)

Further uptake studies for mQD:DOTMA conjugates was done both in SUM-159 and RPE-1 cells using concentrations 100μg/mL and 200μg/mL. The uptake of bioconjugates as compared to mQDs when provided alone was less in both the cell lines. This could be due to the highly positive charge on the conjugates and the increase in size, which might lead to a decrease in the uptake. Upon comparing the ratios in SUM-159A cancerous cells, we can observe that in case of 100ug/mL concentration, 1:2 ratio has relatively higher uptake as compared to others as shown in **Figure 7**. A similar trend was observed in case of 200ug/mL where mQD:DOTMA bioconjugate’s fluorescence intensity was lower indicating lower uptake of the molecule in the cytoplasm as shown in **Figure 8**. Comparing the ratios, we found comparably higher uptake of both 1:2 and 1:10 ratios.

**Figure 7:**
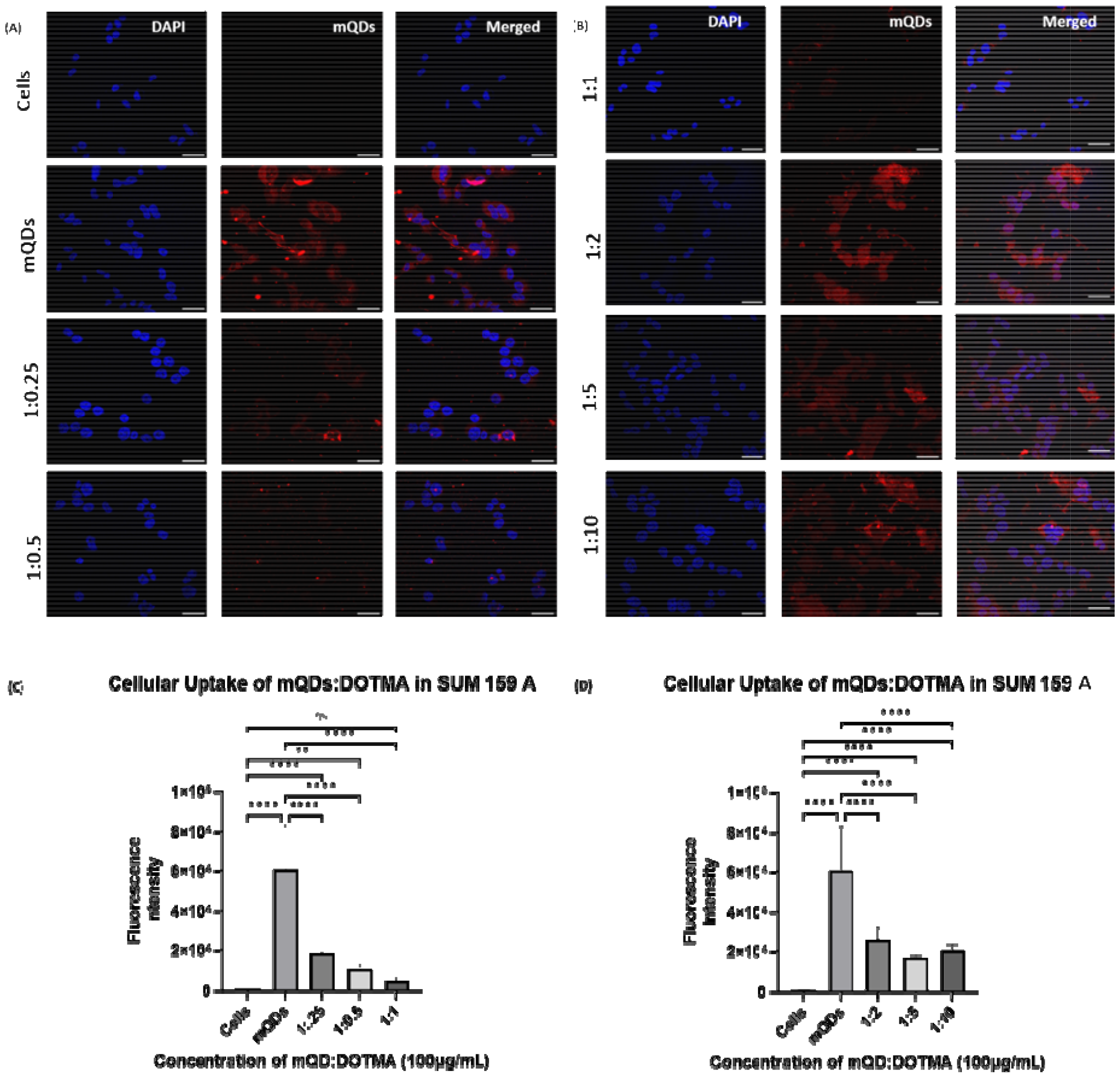
Cellular uptake of mQD:DOTMA conjugates in SUM-159A cells. Concentration - 100μg/mL, Scale bar-5μm (A) Uptake of mQDs and mQD:DOTMA conjugate (1:0.25 and 1:0.5). (B) Uptake of mQD:DOTMA conjugate (1:1, 1:2, 1:5, 1:10). (C) Quantified fluorescence intensity of mQDs and mQD:DOTMA conjugates (1:0.25, 1:0.5, 1:1). (D) Quantified fluorescence intensity of mQD:DOTMA conjugates (1:2, 1:5, 1:10). The statistical significance was tested by one-way ANOVA in the Prism Software and is represented as **** when p < 0.0001 and ns when there is no significant difference. (n = 30 cells per condition)

**Figure 8:**
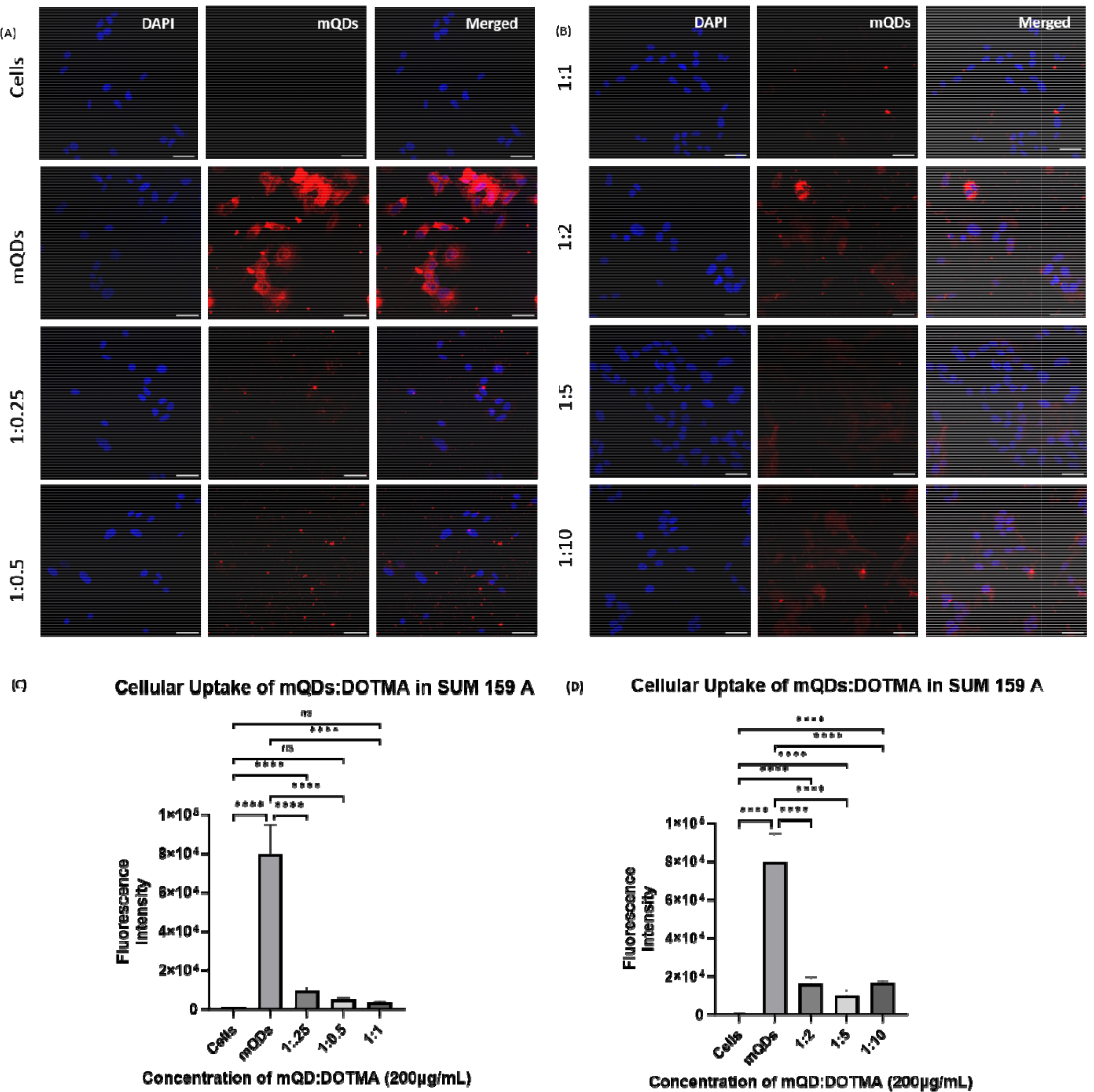
Cellular uptake of mQD:DOTMA conjugates in SUM-159A cells. Concentration - 200μg/mL, Scale bar-5μm (A) Uptake of mQDs and mQD:DOTMA conjugate (1:0.25 and 1:0.5). (B) Uptake of mQD:DOTMA conjugate (1:2, 1:5, 1:10). (C) Quantified fluorescence intensity of mQDs and mQD:DOTMA conjugates (1:0.25, 1:0.5, 1:1). (D) Quantified fluorescence intensity of mQD:DOTMA conjugates (1:2, 1:5, 1:10). The statistical significance was tested by one-way ANOVA in the Prism Software and is represented as **** when p < 0.0001 and ns when there is no significant difference. (n = 30 cells per condition)

In the case of non-cancerous epithelial cells, RPE-1, the uptake of mQD:DOTMA conjugate is relatively lower than only mQDs which can be due to the increase in size and positive charge on the bioconjugate. When the cells are treated with 100μg/mL concentration of mQD:DOTMA, there was an increase in fluorescence intensity in 1:5 and 1:10 which indicates an increase in cellular uptake as shown in **Figure 9**. And when the cells were treated with 200μg/mL concentration of mQD:DOTMA, similar results were obtained **(Figure 10)**.

**Figure 9:**
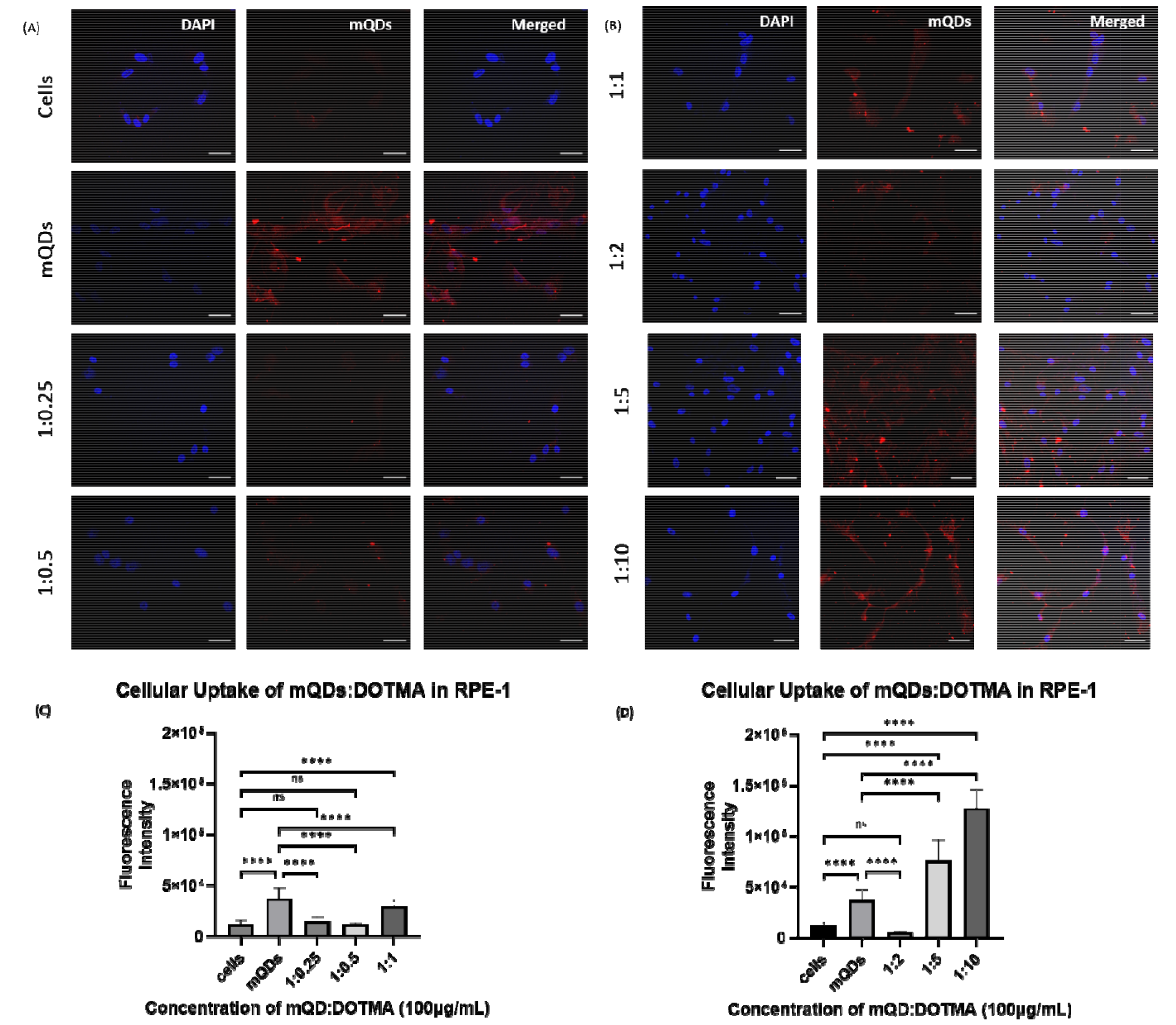
Cellular uptake of mQD:DOTMA conjugates in RPE-1 cells. Concentration - 100μg/mL, Scale bar-5μm (A) Uptake of mQDs and mQD:DOTMA conjugate (1:0.25 and 1:0.5). (B) Uptake of mQD:DOTMA conjugate (1:2, 1:5, 1:10). (C) Quantified fluorescence intensity of mQDs and mQD:DO MA conjugates (1:0.25, 1:0.5, 1:1). (D) Quantified fluorescence intensity of mQD:DOTMA conjugates (1:2, 1:5, 1:10). The statistical significance was tested by one-way ANOVA in the Prism Software and is represented as **** when p < 0.0001 and ns when there is no significant difference. (n = 30 cells per conidition)

**Figure 10:**
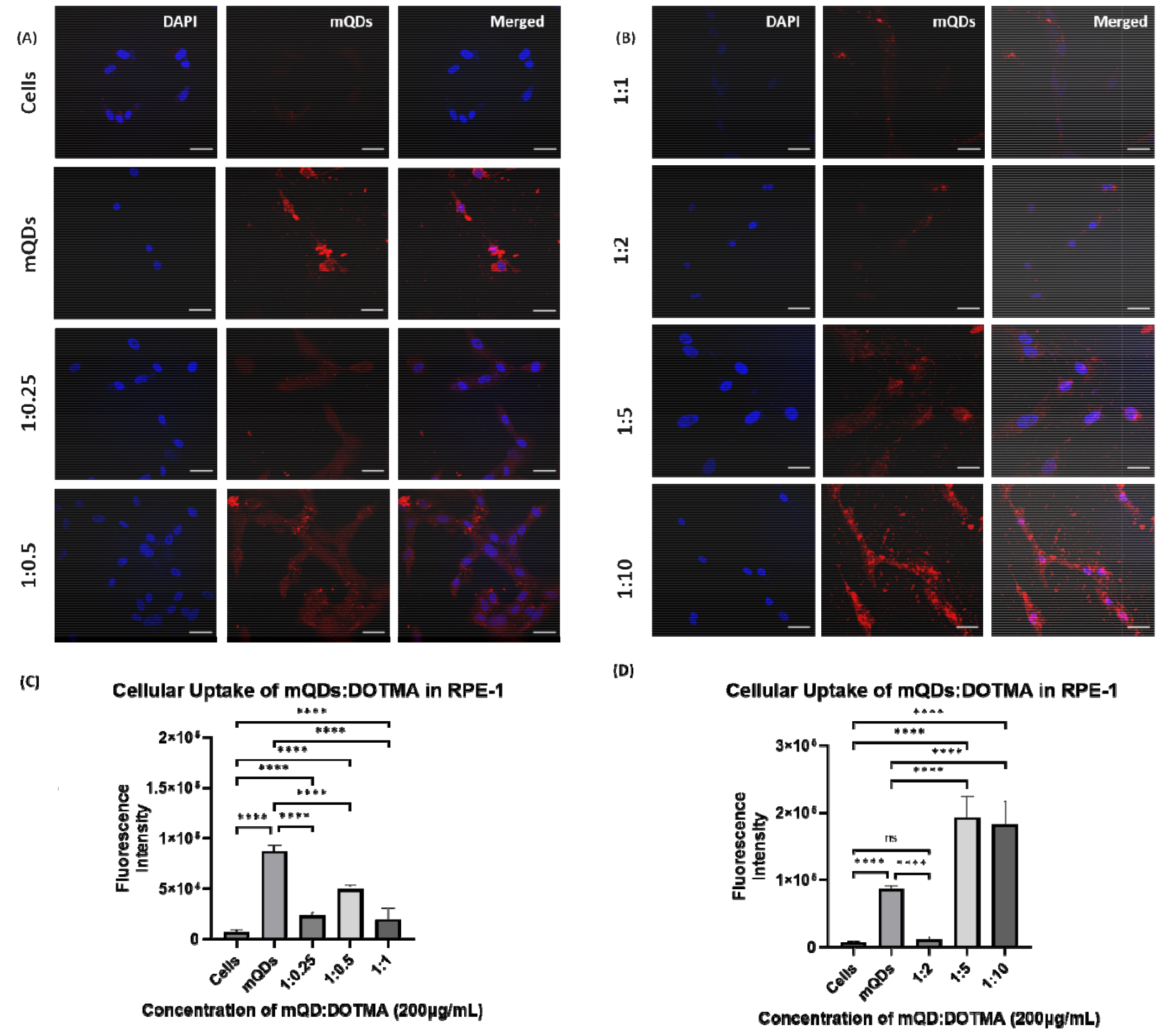
Cellular uptake of mQD:DOTMA conjugates in RPE-1 cells. Concentration - 200μg/mL, Scale bar-5μm (A) Uptake of mQDs and mQD:DOTMA conjugate (1:0.25 and 1:0.5). (B) Uptake of mQD:DOTMA conjugate (1:2, 1:5, 1:10). (C) Quantified fluorescence intensity of mQDs and mQD:DOTMA conjugates (1:0.25, 1:0.5, 1:1). (D) Quantified fluorescence intensity of mQD:DOTMA conjugates (1:2, 1:5, 1:10). The statistical significance was tested by one-way ANOVA in the Prism Software and is represented as **** when p < 0.0001 and ns when there is no significant difference. (n = 30 cells per condition)

## 4. Conclusions and Discussions

In this study, we have successfully synthesized red-emitting carbon quantum dots (mQDs) via a green synthesis approach, offering a sustainable and environmentally friendly method for producing versatile fluorescent probes **(Figure 2)**. These negatively charged mQDs were subsequently conjugated with a small cationic lipid molecule, DOTMA, through electrostatic interaction, resulting in the formation of bioconjugates with various mQD:DOTMA ratios-1:0.25, 1:0.5, 1:0.75, 1:1, 1:2, 1:5, 1:10. Later, characterization analyses were carried out to confirm the conjugation and understand the properties of the biconjugate, including Dynamic light scattering for size and zeta potential, UV-Vis spectroscopy, FTIR, fluorescence spectroscopy, and Atomic Force microscopy (AFM), revealed that the conjugation of mQDs with DOTMA **(Figure 3 and 4)**. The biconjugate thus formed showed a significant increase in fluorescence intensity and photostability compared to standalone mQDs **(Figure 3B and Supplementary figure 1)**. The AFM images also confirmed the successful conjugation of DOTMA with mQDs, showing distinct morphology changes-ring like formation consistent with the formation of the bioconjugate **(Figure 3 F)**.

Next, we studied the effect of this biconjugate *in-vitro* in cancerous cells (SUM-159A) and epithelial cells (RPE-1). Interestingly, our cellular uptake studies using fluorescence microscopy demonstrated that the uptake of the mQD:DOTMA bioconjugate in SUM-159 (cancerous cells) was lower than that of mQDs alone **Figure 7 and 8**. This reduced uptake can be attributed to the increased size and zeta potential of the bioconjugates, which may hinder their cellular internalization^21^**(Figure 11)**. However, specific ratios of mQD:DOTMA bioconjugates showed higher uptake in RPE-1 cells, suggesting a potential dose-dependent effect on cellular internalization. Upon comparing different ratios of mQD:DOTMA, we found out that in SUM-159A, the 1:2 and 1:10 ratio exhibited relatively higher uptake compared to others at both the concentration of 100 and 200 μg/mL, as depicted in **Figure 7**. Similarly, for RPE-1 the fluorescence intensity of the mQD:DOTMA conjugate of ratio 1:5 and 1:10 is higher than mQDs at both the concentrations-100 and 200μg/mL, indicating increased cytoplasmic uptake, as illustrated in **Figure 9 and 10**.

**Figure 11:**
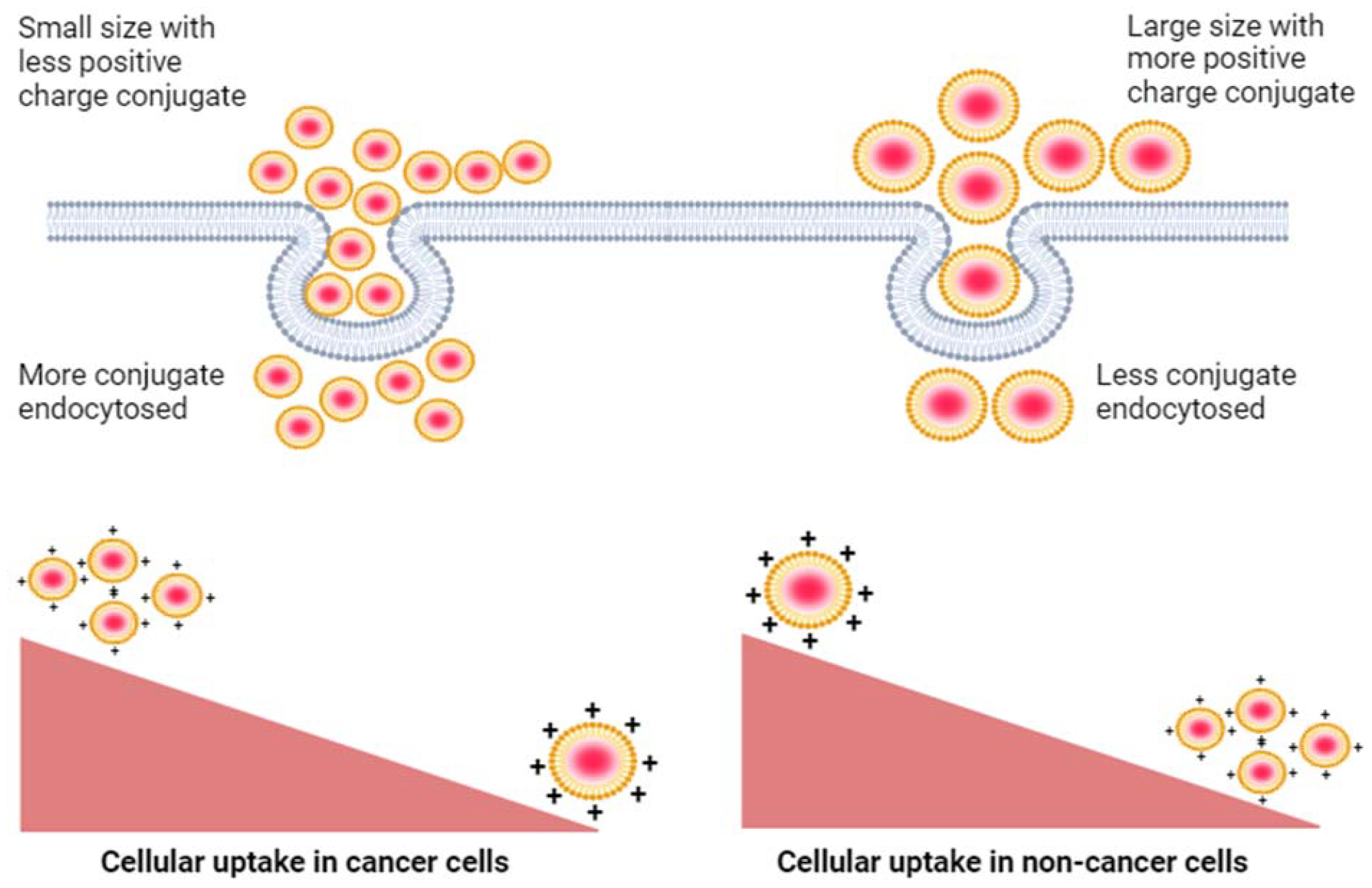
Decreased cellular uptake of mQD-DOTMA bioconjugate in cancer cells as compared to non-cancer cells due to increased size and zeta potential of the bioconjugate

Furthermore, cytotoxicity assays revealed that while both mQDs and DOTMA alone exhibited toxicity across all tested concentrations (100, 200, 300μg/mL) in SUM-159A and RPE-1 cells^22^, some ratios of the mQD:DOTMA bioconjugate showed quenching of cytotoxicity of individual molecules and maintained high cell viability. In SUM-159A 1:0.5 at concentration 200 and 300μg/mL showed 104% and 104% respectively. In RPE cells these bioconjugate have increased the cell viability. The maximum cell viability was observed in 1:2 at 100μg/mL (cell viability 227%), 1:2 at 200μg/mL (cell viability 315%) and 1:1 at 300μg/mL (cell viability 370%) This indicates the potential of these bioconjugates to mitigate cytotoxic effects which can be further explored in bioimaging and potential therapeutics. **(Figure 5)**.

The versatility of mQD:DOTMA bioconjugates opens numerous possibilities for biomedical applications, including targeted drug delivery, cellular imaging, and therapeutics. The enhanced fluorescence intensity and photostability make them ideal candidates for sensitive and prolonged imaging of cellular processes. Furthermore, the ability to modulate cytotoxicity through ratio optimization underscores their potential for safe and effective drug delivery systems.

In conclusion, the mQD:DOTMA bioconjugates represent a promising platform for a wide range of biomedical applications, offering enhanced fluorescence properties, tunable cytotoxicity, and potential therapeutic benefits. Future studies should focus on further optimizing the bioconjugate ratios, investigating their efficacy in vivo, and exploring additional characterization techniques to fully realize their translational potential in biomedical research and clinical practice. This shows that the bioconjugate is not toxic and can be used as therapeutics.

## Acknowledgement

The authors sincerely thank all the members of the D.B. group for critically reading the manuscript and for their valuable feedback. P.Y. thanks IITGN-MoES, GoI, Ph.D. fellowship; P.Y. acknowledges Director’s fellowship from IITGN for additional fellowship. D.B. thanks SERB, GoI for Core Research Grant, STARS-MoES for research grant, DST-Nidhi Prayas for the start-up grant, and Gujcost-DST, GSBTM, BRNS-BARC, and HEFA-GoI for research grants. D.B. is a member of the Indian National Young Academy of Sciences (INYAS). CIF at IIT Gandhinagar for confocal and AFM facility, CRTDH lab in chemical engineering department at IITGN for fluorescence studies, DLS and FTIR facility in material science department at IITGN are acknowledged.

## Notes

### Competing Interest Statement

The authors have declared no competing interest.

