## Supplementary material for "Cellular uptake and viability switch in the properties of lipid-coated carbon quantum dots for potential bioimaging and therapeutics": Supplimentary file

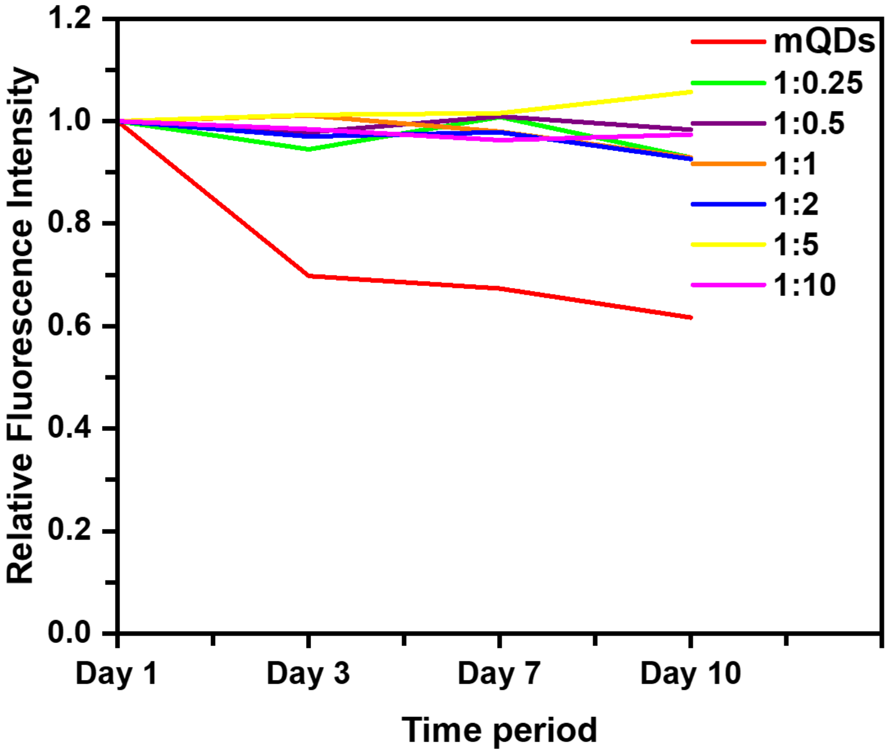


**Figure S1:** The relative fluorescence intensity was measured for 10 days. It shows that the fluorescence intensity of mQDs has dropped to about 61.8% than on day 1. Whereas in case of mQDs coated with DOTMA (1:0.25, 1:0.5, 1:1, 1:2, 1:5, 1:10) we observed that the fluorescence intensity was 93%, 98%, 93%, 93%, 106%, 97%. We observed that in ratio 1:5 the fluorescence is increasing with time.
